# Behavioral and anatomical effects of complex rearing after day 7 neonatal frontal lesions vary with sex

**DOI:** 10.1101/2024.07.08.602597

**Authors:** Robbin Gibb, Bryan Kolb

**Author notes:** **Correspondence address:** Bryan Kolb, Canadian Centre for Behavioural Neuroscience, University of Lethbridge, Lethbridge, AB, Canada, T1K 3M4, FX.

## Abstract

Rats received medial frontal lesions, or sham surgery, on postnatal day 7 (P7), and were placed in complex environments or standard lab housing at weaning. Three months later the animals were tested in a spatial learning task and in a skilled reaching task. Rats with P7 lesions had smaller deficits in spatial learning than similar adult operates, and males performed better than females. Complex housing further improved the performance of male, but not female, lesion rats. The opposite was seen in motor performance as female lesion rats performed better than males and benefited from experience whereas males did not. There was a correlation between behavior and dendritic change as complex housing reversed lesion-induced dendritic atrophy, an effect that was greater in males. P7 frontal males, but not females, showed an increase in spine density that was reversed by complex housing. Experience thus can affect functional and anatomical outcome after early brain injury, but the effects vary with sex.

---

Pediatric brain insult, whether it be vascular or traumatic, is a leading cause of long-term disability in children with injuries as infants or toddlers (e.g., Arambula et al., 2019; Sullivan et al., 2023). Little is known about effective treatments, however, and there are contradictory findings related to possible sex differences in long-term outcomes in children, and whether possible treatments might vary with sex. With these questions in mind, we elected to use our animal model to investigate complex housing as a possible treatment, and to consider the role of sex in both behavioral and anatomical outcomes.

Damage to the cerebral cortex of the rat produces an age-related functional outcome in which damage in the first few days of life produces much larger chronic functional deficits than similar injuries at days 7-10 (e.g., Kolb, 1995; Kolb & Whishaw, 1989). The age-related difference in functional recovery is presumed to be related to age-related differences in ongoing processes of brain development in the first and second weeks (e.g., Kolb, 1995; Nonneman et al., 1983; Rice & Baronne, 2000; Schneider & Koch, 2005). What is not known, however, is whether treatments that influence recovery after injury are equally effective at the two age points. In particular, whereas there are several studies showing that complex housing (also referred to as enriched experience) enhances recovery after cortical injury in the first few days of life (e.g., Kolb and Elliott, 1987), virtually nothing is known about the effects of such experience on injuries from days 7-10, a time at which the effects of the injuries are much attenuated relative to the effects resulting from lesions in the first few days of life. Postnatal day 7-10 is arguably more relevant to human children with perinatal brain injuries because it would be equivalent to postnatal injury rather than prenatal equivalent days 1-5. It could be hypothesized that the brain is more plastic after day 7-10 lesions so we would predict a large effect of experience. But, it could also be proposed that there are limits to plasticity after early injury and little additional benefit would accrue from the complex housing.

We therefore chose to examine the effects of complex housing on rats with medial frontal lesions on day 7. Because we had found previously that the behavioral and morphological effects of lesions at this age were sexually-dimorphic (Kolb & Stewart, 1995), we also wondered if the effect of rearing experience might also show sex-related differences, especially in the lesion animals. Our prediction was that because females show less spontaneous recovery than males, they might show more benefit from the complex housing therapy.

We chose to examine behavior on a cognitive task, the Morris water task (Morris, 1981), as well as a motor task, the Whishaw reaching task (Whishaw et al., 1991). The Morris task is sensitive to the effects of medial prefrontal lesions (Sutherland, Kolb & Whishaw, 1982) and has proven to be a reliable measure of the effects of behavioral and pharmacological treatments on functional recovery after early medial frontal lesions (Kolb & Elliott, 1987; Kolb & Sutherland, 1992; Kolb, Sutherland & Whishaw, 1983). Skilled reaching is profoundly disrupted by medial prefrontal injuries at any age (e.g., Kolb, Petrie & Cioe, 1996; Whishaw et al., 1991) and shows less spontaneous recovery than direct motor cortex injury that spares the medial prefrontal region. The question is whether complex housing could improve motor functions in rats with P7 lesions.

We measured dendritic arborization in parietal and occipital cortex because both regions show increased dendritic arborization in response to complex housing in male rats (Kolb, Gibb & Gorny, 2003), and because changes spine density in parietal cortex are correlated with functional outcome in male (Kolb, Stewart & Sutherland, 1997), but not in female rats (Kolb & Stewart, 1995). We were unaware of previous studies on the effects of early lesions or experience in lesion animals in dendritic organization on posterior cortical neurons in rats, but we felt it was important to know how widespread the effects of the day 7 lesions might be on cortical organization. We had shown previously that the entire cortical mantle was thinner than normal after day 7 lesions so it seemed likely that we would find dendritic atrophy in the occipital cortex.

Although it would be useful to measure the effects of the injuries, experience, and training on motor cortex, we chose not to do this because we did not have animals without training and thus we could not distinguish the selective effects of either training or experience on the motor cortex neurons, which are known to show increased dendritic arbor with reach training (e.g., Kolb et al., 2008; Withers & Greenough, 1988). Parietal cortex neurons do not normally show reaching-related changes in dendritic organization, however (B. Kolb, unpublished observations). In addition, the early injuries disrupt normal lamination in the adjacent motor cortex and shift the location of the motor maps making it difficult to determine whether cells in the lesion and control animals were in the same subregion (Williams et al., 2006).

## MATERIALS AND METHODS

### Subjects

The study was done with 40 Long-Evans rats derived from 5 litters of Charles-River strains, which were divided into four groups of ten rats using a cross litter design (5M and 5F per group): lab control, lab frontal, enriched control, and enriched frontal. The rats in each age group were assigned to either the lab or enriched housing such that body weight was approximately equal in the lab and enriched groups and that approximately equal numbers of animals in each of the lesion and treatment groups came from the same litter.

### Surgical procedures

All animals were anesthetized on the seventh day after birth by cooling them in a Thermatron cooling chamber until their rectal body temperatures were in the range of 18-20°C. For the frontal rats the frontal bone was removed by cutting it with iris scissors, and frontal decortication was achieved by gentle aspiration. The intent was to remove the medial subfields of the prefrontal cortex including the presumptive Zilles’ (1985) regions Cg 1, Cg 3, and PL as well as the medial portion of Fr 1 of the motor cortex. As soon as medial frontal decortication was achieved, the animals’ scalps were sutured with silk thread. The normal control group animals were anesthetized in the same manner, and the skin was incised and sutured.

### Complex housing procedures

The rats were reared with their mothers and after weaning at day 22 they all were housed either in 65 X 26 X 18 cm stainless steel hanging cages (3-4 per cage) or placed into the complex housing conditions for 3 mo. The complex housing took place in large vertically-organized pens measuring 63 X 148 X 187 cm. Three of the walls (sides and front) were made of hardware cloth. The back wall was made of plywood covered with blue Arborite, as was the ceiling and floor. Two stainless steel cages (22 X 26 X 18 cm) were attached to the upper part of the front wall, and another was placed on its side on the cage floor. There also were runways attached to the back wall, which allowed animals to navigate from the floor to a shelf near the top without having to run up the hardware cloth walls. There were two similar pens with 4-6 same-sex animals housed in each, with roughly equal numbers of lesion and sham animals in each pen. The pen floor was covered with about 10 cm of sawdust bedding. The pen was filled with toy objects (plastic pipes, children’s toys, branches, etc. The objects were changed weekly. The animals were left undisturbed except for daily feeding and weekly cleaning of the pen.

### Behavioral assessment

#### Morris Water Task

The method used in this test was similar to that described elsewhere (Sutherland et al., 1983) and is based on the original task described by Morris (1980). The maze was a circular pool (1.5 m diameter x 0.5 m deep) with smooth white walls. The pool was filled with approximately 20°C water, mixed with 1 L of skim milk powder or just enough to render the water opaque. A clear Plexiglas platform (11 x 12 cm) was placed in a constant position inside the pool approximately 12 cm from the wall. The water level was adjusted so that the platform stood 2 cm below the surface of the water. The platform was invisible to a viewer outside the pool and to a rat swimming in the water. A trial consisted of placing a rat into the water at one of four locations (north, south, east, or west) around the pool’s perimeter. Within a block of four trials each rat started at the four locations in a random sequence, and each rat was tested for eight trials a day over five consecutive days, for a total of 10 trial blocks. If on a particular trial a rat found the platform, it was permitted to remain on it for 10 seconds. A trial was terminated if the rat failed to find the platform after 90 seconds. Each rat was returned to its holding cage for approximately 5 minutes before the next trial commenced. The swimming path for each rat on every trial was traced by the experimenter and latency to find the platform was recorded. The heading error was calculated manually from the drawings.

### Skilled Reaching

The animals were tested in a skilled reaching task after the completion of the water task. The reaching procedure, developed by Whishaw et al. (1991), was used to assess the skilled forelimb movements of each rat after being trained to reach for chicken feed pellets in Plexiglas cages (28 cm deep x 20 cm wide x 25 cm high). The front and floor of each cage were constructed with 2 mm bars separated from each other by 1 cm, edge to edge. A tray (5 cm deep x 2 cm wide x 1 cm high) containing chicken feed pellets, was mounted in front of each cage. To obtain food, the rats had to extend the forelimb through the bars, grasp, and retract the food pellet. The food tray was mounted on runners to adjust the distance of the food from the bars. Distance adjustments ensured that each rat could not simply rake the food into the cage. Bars on the floor ensured that if the rat dropped the pellet, it would irretrievably lose it and would have to reach again. Rats were trained on the task for a maximum of three weeks before video-taping. During the first week, the rats were grouped in pairs in the reaching cages for one hour a day to allow the rats to adapt to their new surroundings. The food deprivation schedule commenced during the first week, and each rat was provided with 15 grams of laboratory rodent food daily following the training period. The rats were subsequently trained individually for one hour each day during the second week, whereas during the third week, this training period was shortened to 5–15 minutes a day. Five minutes of continuous reaching activity for each rat was videotaped and scored when the rats were approximately 5 months of age. If the rat made a reaching movement (forepaw inserted through the bars, but no food was grasped or it was dropped), it was scored as a “reach”. If the rat obtained a piece of food and consumed it, the movement was scored as a “reach” and a “hit.” Scoring was achieved by calculating the percentage of hits to total reaches for each animal’s preferred forelimb. Left and right paw reaches and hits were recorded separately. Most rats had a strong preference for one paw over the other, and many only reached with one paw, so we used the reaching from the preferred paw (>60% of reaches with one paw) as the measure of reaching ability. In all cases this paw was more accurate. In animals showing no clear paw preference we used the data from the more accurate paw as the measure.

### Anatomical Methods

Following the conclusion of the enriched housing the animals were given an overdose of sodium pentobarbital and intracardially perfused with 0.9% saline. The brains were removed and weighed before being immersed whole in 20 ml of Golgi-Cox solution. The brains were left in the solution for 14 days before being placed in a 30% sucrose solution for 2-5 days, Golgi-fixed brains were left in solution for 14 days before being placed in a 30% sucrose solution for 2 days, cut on a Vibratome at 200 µm, and processed using a procedure described in Gibb & Kolb (1998).

Layer III pyramidal cells in Zilles’ areas Oc1 or Layer III cells in Zilles’ area Par1 were traced using a camera lucida at 250X. To be included in the data analysis, the dendritic trees of pyramidal cells had to fulfill the following criteria: (a) the cell had to be well impregnated and not obscured with blood vessels, astrocytes, or heavy clusters of dendrites from other cells; (b) the apical and basilar arborizations had to appear to be largely intact and visible in the plane of section. Dendritic length was analyzed by using the method of Sholl (1956). The number of intersections of dendrites with a series of concentric spheres at 25 µm intervals from the center of the cell body was counted for each cell. Statistical analyses were performed by averaging across ten cells per hemisphere. An estimate of mean total dendritic length (in µm) can be made by multiplying the mean total number of intersections by 25. For branch order analysis, each branch segment was counted and summarized according to methods of Coleman and Riesen (1968): branches emerging from either the cell body (basilar) or the primary apical dendrite (apical) were first order. After the first bifurcation, branches were considered second order, etc. Quantification of each branch type using this method provided an indication of dendritic arbor complexity.

Spine density was measured in the occipital cortex from one apical dendritic branch in the terminal tuft and one basilar terminal branch. Spine density measures were made from a segment greater than 10 µm in length, and usually about 50 µm. The dendrite was traced (1000X) using a camera lucida drawing tube and the exact length of the dendritic segment calculated by placing a thread along the drawing and then measuring the thread length. Spine density was expressed as the number of spines per 10 µm. No attempt was made to correct for spines hidden beneath or above the dendritic segment, so the spine density values are likely to underestimate the actual density of the dendritic spines. Spine density was measured on the terminal tip and one oblique branch (second order branch that did not reach layer I) branch on the apical dendrites and one terminal tip on the basilar dendrites. Two measures of dendritic spines were made on the apical branches because there was an obvious difference in density between the oblique and terminal branches. There was no such difference visible in the basilar branches.

## BEHAVIORAL RESULTS

### General Behavioral Observations

As in our previous studies, the enriched animals were frequently observed interacting with the objects and moving about the cages. The complex housing pen allowed the animals to engage in considerable vertical activity and the animals took obvious advantage of this opportunity. In fact, they became extremely agile and if offered food treats anywhere along the hardware cloth walls, they could rapidly run up and down the runways and walls. One unexpected observation was that given a choice, the rats in pens preferred to sleep in cages that were hung a meter or more off the floor. They were rarely observed sleeping on the cage floor. If nesting materials such as paper towel were available on the pen floor both male and female animals would carry it up to the hanging cages where they built nests. Because both control and frontal rats were housed together, it was not possible to tell if frontal rats carried bedding or made nests but in view of our previous findings that frontal animals do not hoard or nest build, it seems unlikely that they did (e.g., Kolb and Whishaw, 1983).

### Morris Water Task

#### General observations

Control rats quickly learn that there is a hidden platform in the tank and rapidly learn to swim to the platform from any start location. By the third trial block the control rats asymptoted with an escape latency of about 5 sec. Frontal rats typically take longer to leave the wall and begin to search, leading to much higher latencies on the first trial block. As in our previous experiments, we found only a small effect of complex rearing on the water task performance of control animals (e.g., Kolb and Gibb, 1990; Kolb and Elliott, 1987). Although rats with P7 lesions showed significant recovery on this task relative to rats in our previous studies with similar removals in the first few days of life or in adulthood (e.g., Kolb, 1987), they were still impaired relative to sham-operated controls, confirming a result that we have found previously (e.g. Kolb, 1987). Furthermore, as in an earlier study (Kolb and Stewart, 1995), we found a sex-related difference as females with P7 lesions were more impaired than males with similar lesions (Figs 1, 2). Finally, even though female frontals had larger deficits than males, they did not show any significant benefit from the complex housing whereas the males did.

**Figure 1.**
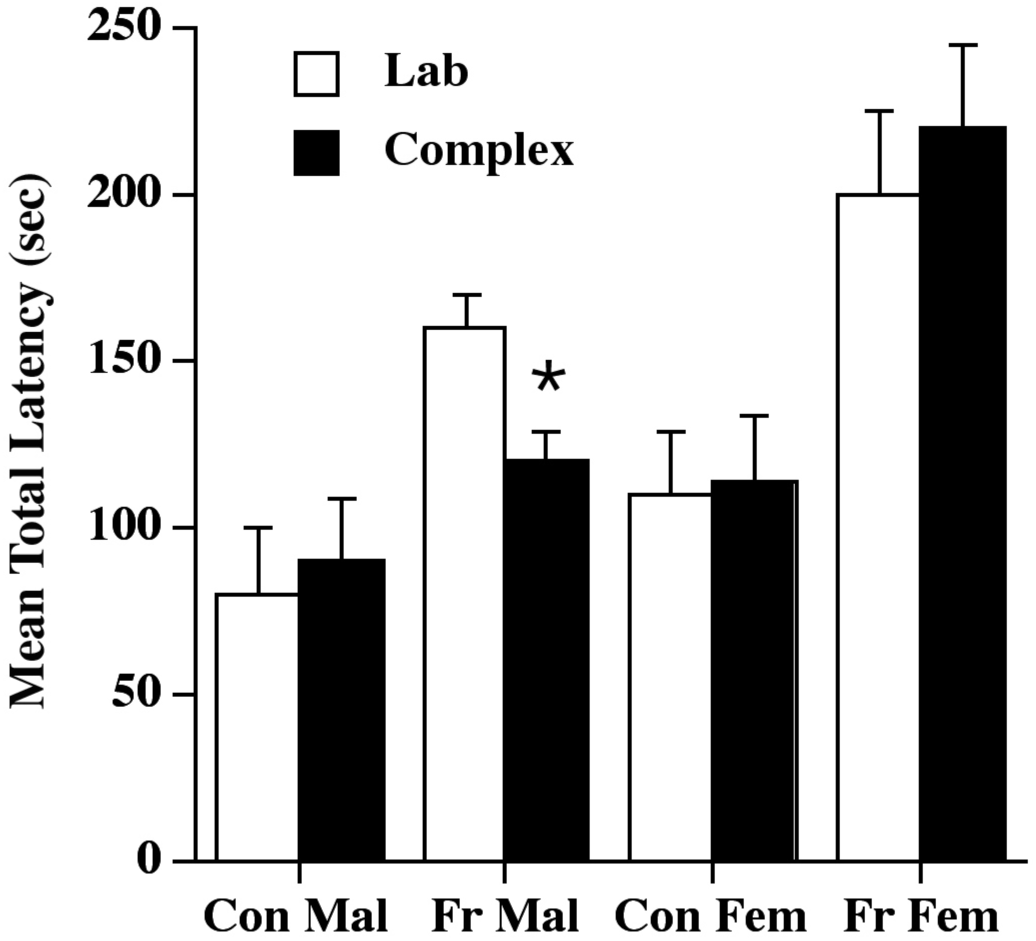
Summary of Morris water task performance of the P7 frontal operates who were placed in complex environments (or standard lab housing) at weaning. Males with frontal lesions showed smaller deficits than females but only the male frontal group benefited from the complex housing experience. Abbreviations: Con Mal=control male; Fr Mal=frontal male; Con F=control female; Fr Fem=frontal female; R=reversal of the platform location to opposite side of the tank.

A four factor ANOVA (Group X Experience X Sex X Trial Block) was performed across Trial Blocks 1-9, and found a significant main effect of lesion (F(1,32)=5.5, p=.02), sex (1,32)=18.0, p<.0002), and trial block (F(8,256)=73.1, p<.0001) but not of experience (F(1,32)=0.2, p=.90). There were two significant interactions: Lesion X Trial Block (F(8,256)=4.3, p<.001) and Experience by Trial Block (F(8,256)=2.3, p=.02). The remaining interactions were small and nonsignificant, (F’s<1, p’s>.28).

The two significant interactions are instructive. First, the Block X Lesion interaction reflected that the lesion animals learned the task more slowly than the control animals. Second, the Block X Experience interaction reflected that the animals housed in the complex environments learned the task more quickly than the lab-reared animals. Although there was no interaction with Sex, posthoc tests showed that only the male frontals showed an overall benefit from the complex housing as shown in Figure 1.

Finally, to determine if the animals had successfully learned the location of the hidden platform, we compared the performance of Trial Block 9, which was the last set of trials to the original platform location, to Trial Block 10 (marked as ‘R’ in Figure 2), which was a set of four trials to a new location. A four way ANOVA (Group X Experience X Sex X Trial Block 9 and 10) found a significant main effect of the comparison of trial blocks 9 and 10 (F(1,32)=50.8, p<.0001) but no other effects were significant (F’s<1.5, p>.25). Thus, all groups showed a significant disruption in performance from Trial Block 9 to 10, indicating that they knew the original location of the platform.

**Figure 2.**
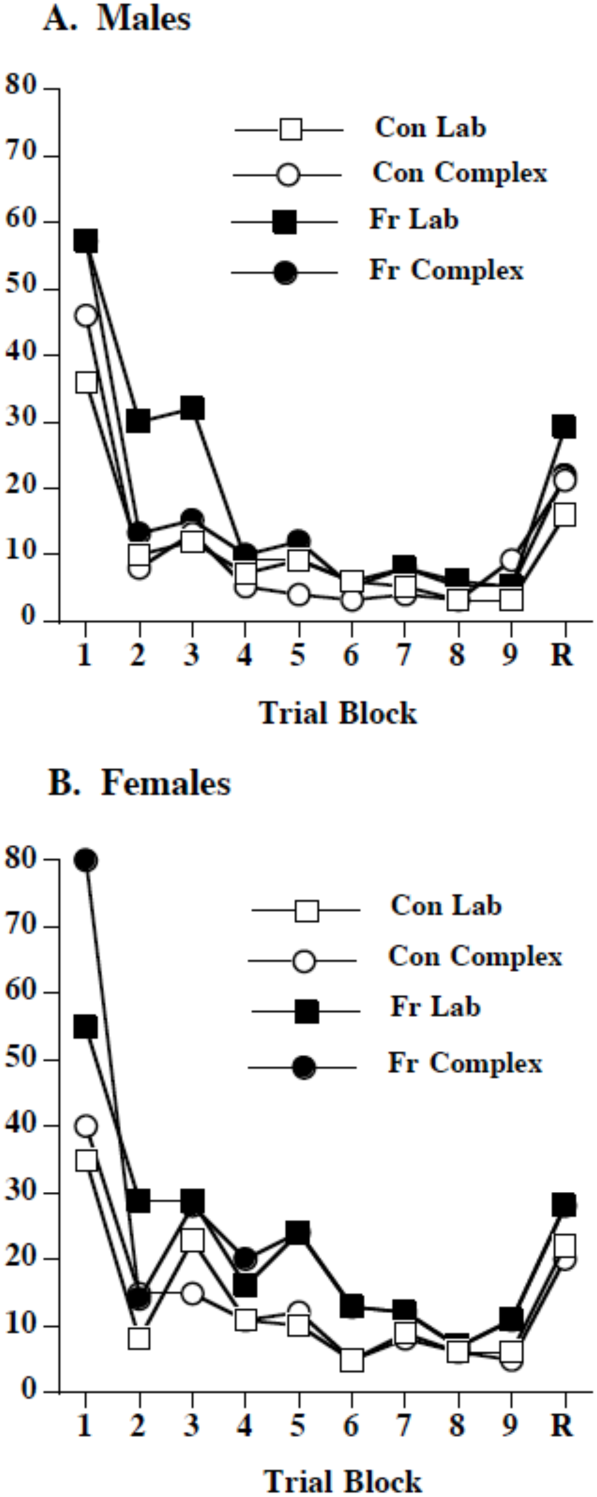
Morris water task performance by trial block. The benefit of complex housing in the males was largely in the second and third trial blocks. Overall, complex-housed frontal males performed as well as controls by the second trial block whereas the females only reached control level on the seventh trial block.

### Skilled Reaching

Control rats acquire the reaching task quickly, reaching asymptotic performance around 60% accuracy. As summarized in Figure 3, rats with frontal lesions in infancy do very poorly at this task with the males performing around 8% and the females around 30% accuracy. The females, but not the males showed a beneficial effect of the complex housing on reaching performance. There was no overall difference in the amount of reaching in the control and frontal animals, the average being about 75 reaches during the filming period.

**Figure 3.**
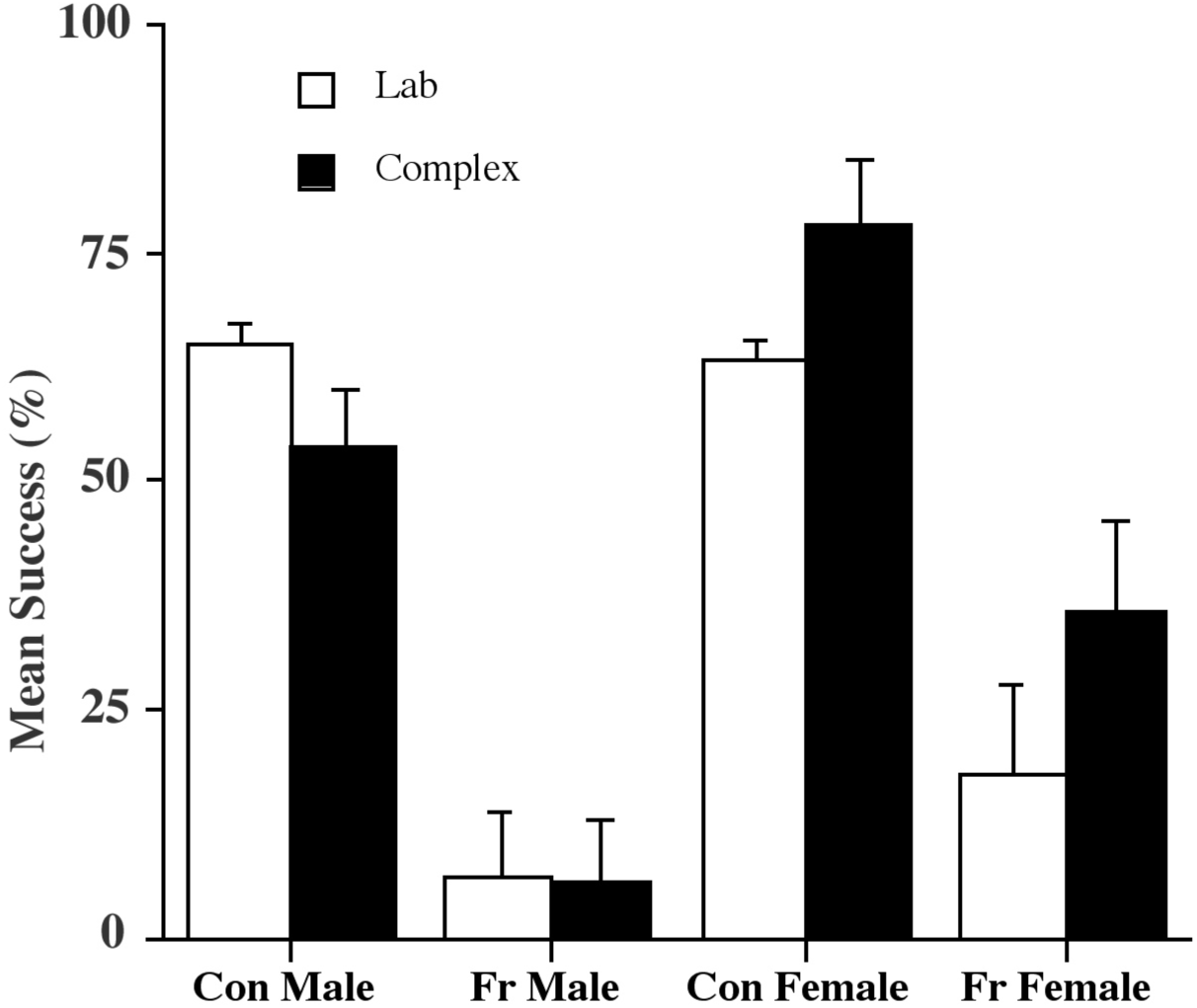
Summary of reaching accuracy in a skilled reaching task. All frontal groups were impaired at the task but females performed better. Only females benefited from complex housing.

A three factor ANOVA (Group X Experience X Sex) revealed main effects of lesion group (F(1,33)=114, p<.0001), sex (1,33)=14.2, p=.001, but not of experience (F(1,33)=1.5, p=0.24. In addition, the Sex X Experience interaction was significant (F(1,33)=7.1, p=.01), but none of the other interactions were significant (F’s<1.5, p’s>.2). The Sex X Experience interaction reflects the obvious sex difference in the effect of experience shown in Figure 3: Females benefited from experience whereas males did not.

## ANATOMICAL RESULTS

### General Observations

Gross inspection of the lesions showed that the lesions were roughly as intended with removal of Zilles’ areas Cg1, Cg3, and the medial portions of Fr1, as well as the more lateral portions of Fr1 in some animals (Figure 4). There were no obvious differences in lesion size between the lab- and complex-housed frontal rats nor were they any obvious sex differences.

**Figure 4.**
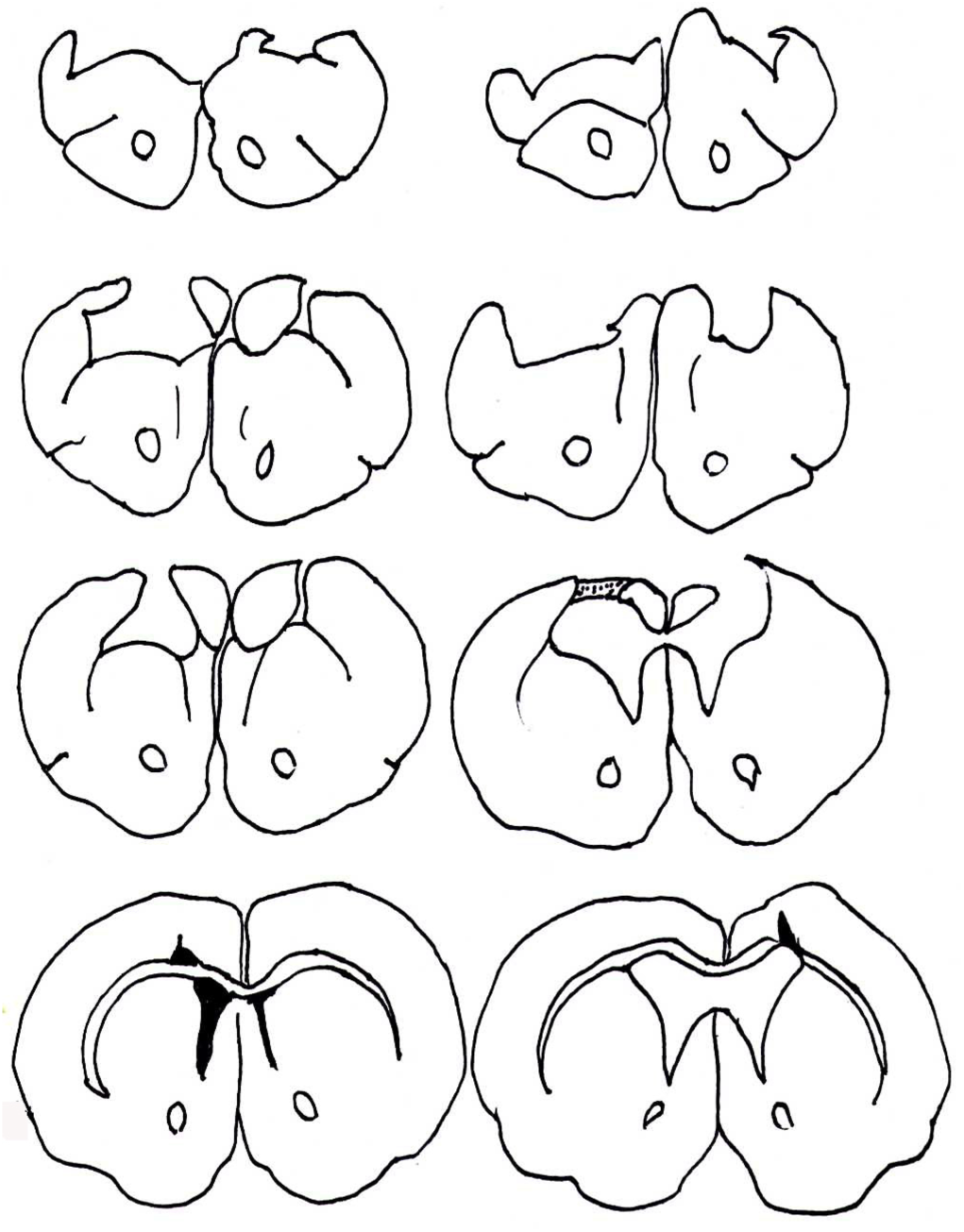
Schematic illustrations of the extent of the frontal lesions of female animals reared in laboratory cages (left) versus complex environments (right). There was no obvious difference in the size of the lesions in males and females (males not shown).

### Dendritic arborization and spine density

#### Dendritic branching

There were three overall effects. First, there was a lesion effect on dendritic branching as frontal animals showed a significant atrophy in dendritic branching in both the parietal and occipital cortex, the effect being larger in males in parietal cortex and larger in females in occipital cortex. Second, as illustrated in Figures 5-8, the effect of complex housing was larger in the males than the females. Third, complex-housing reversed the lesion-induced changes in dendritic arborization, the effect being larger in males than females. Three-way ANOVAs (Lesion x Experience x Sex) were performed separately on the apical and basilar fields for the parietal and occipital regions respectively, and for the dendritic length and branching, respectively.

**Figure 5.**
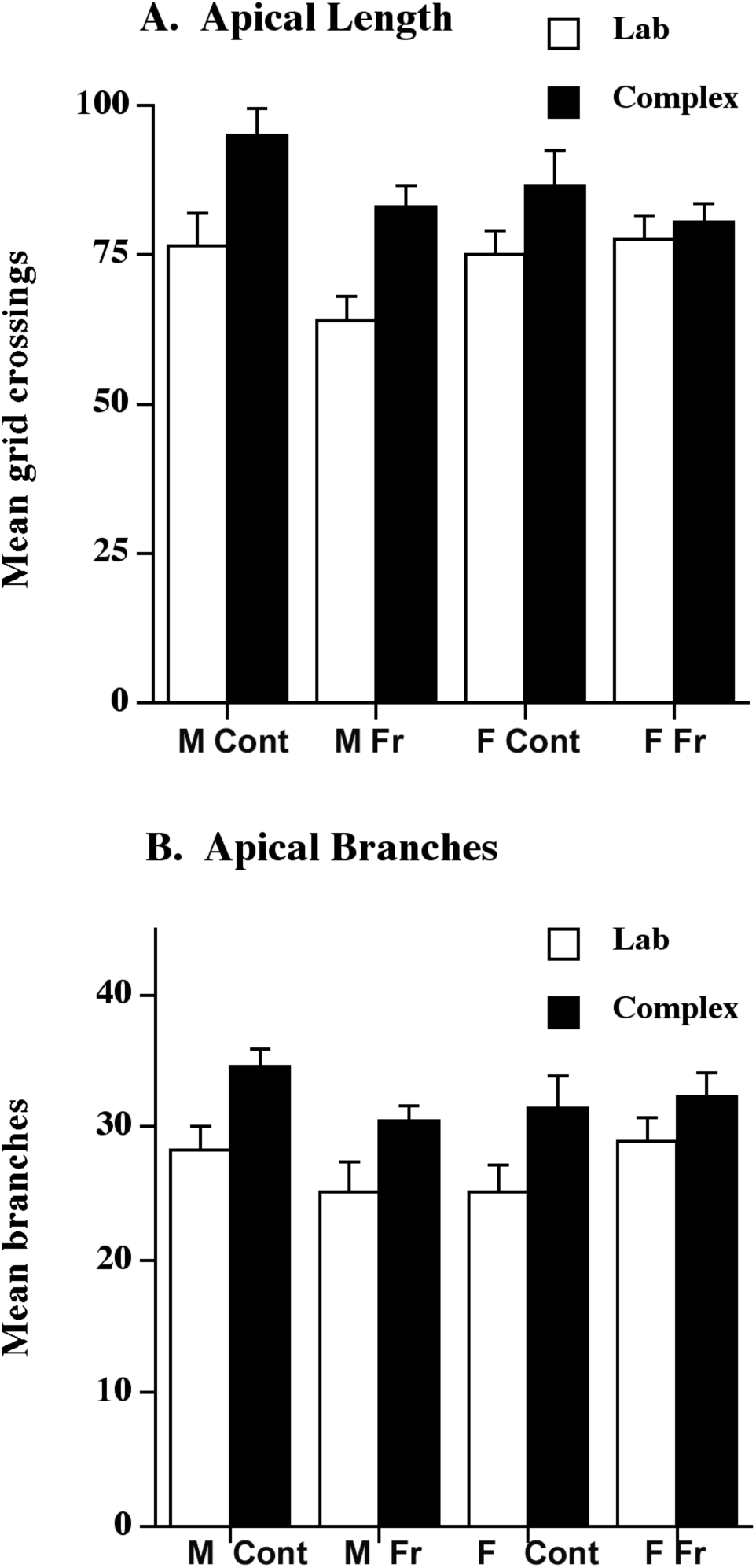
Summary of the dendritic changes seen in the apical length (top) and branching (bottom) of layer III pyramidal neurons in Zilles’ area Par 1. Complex housing increased dendritic length and branching in all groups. P7 frontal lesions produced an atrophy of dendritic arbor in males and complex housing reversed this atrophy.

**Figure 6.**
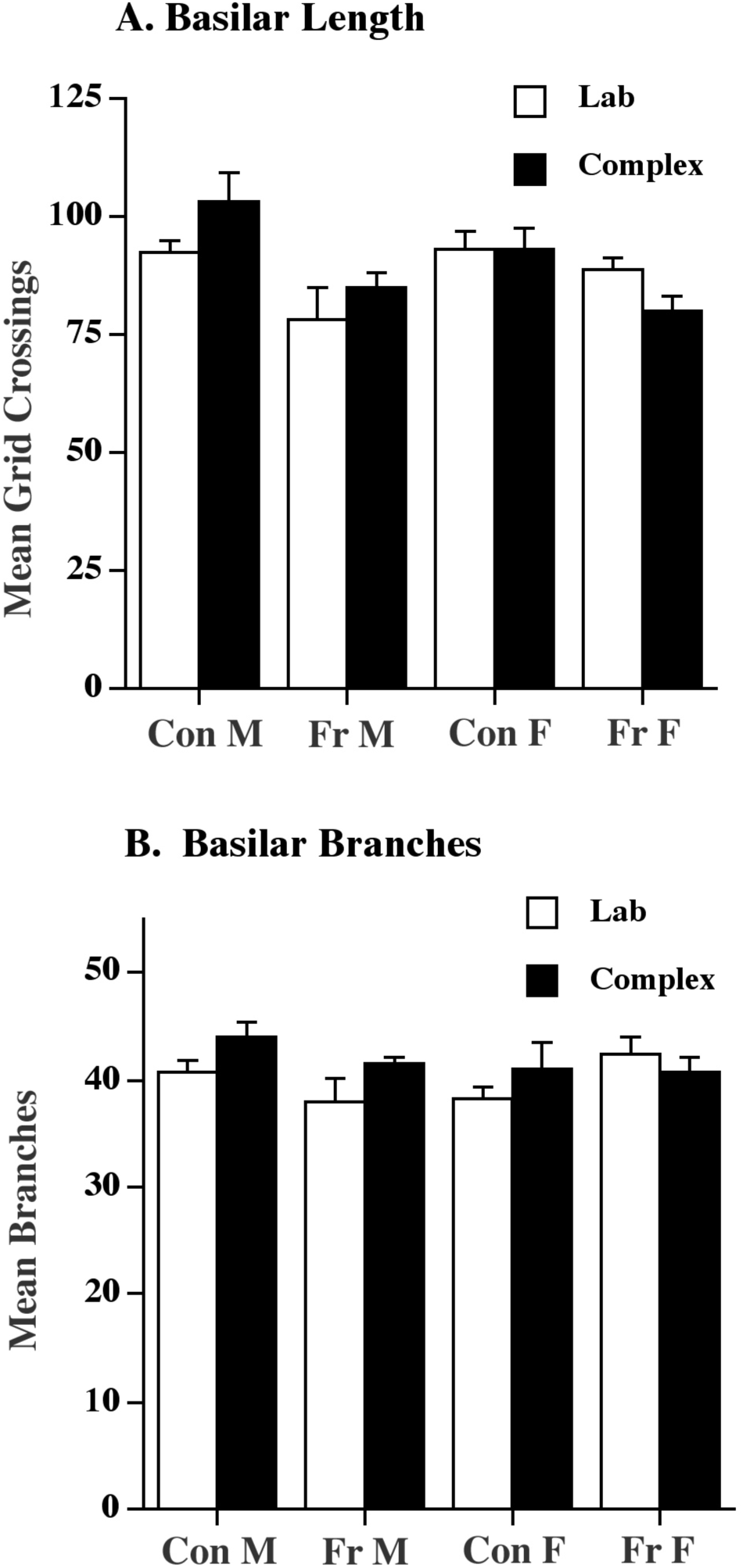
Summary of dendritic changes seen in the basilar length (top) and branching (bottom) of layer III pyramidal neurons in area Par 1. Complex housing increased dendritic length and branching in males but not females.

**Figure 7.**
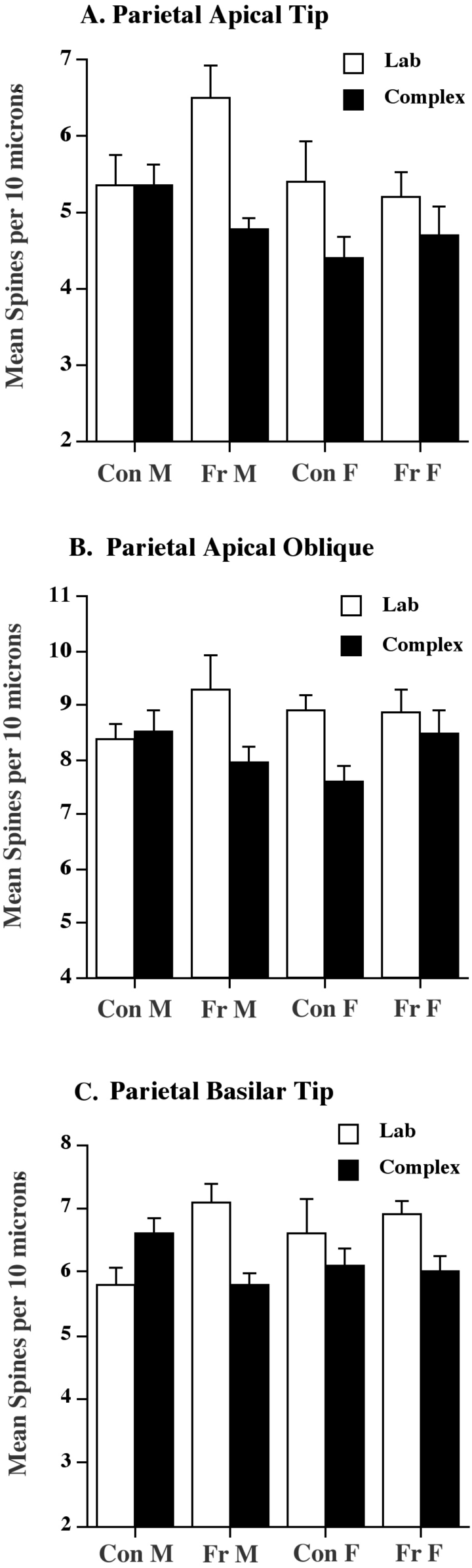
Summary of the changes in spine density in layer III neurons in parietal cortex. Frontal lesions increased spine density in males but not females. Complex housing decreased spine density in the lesion animals in both sexes but did not do so in control males.

**Figure 8.**
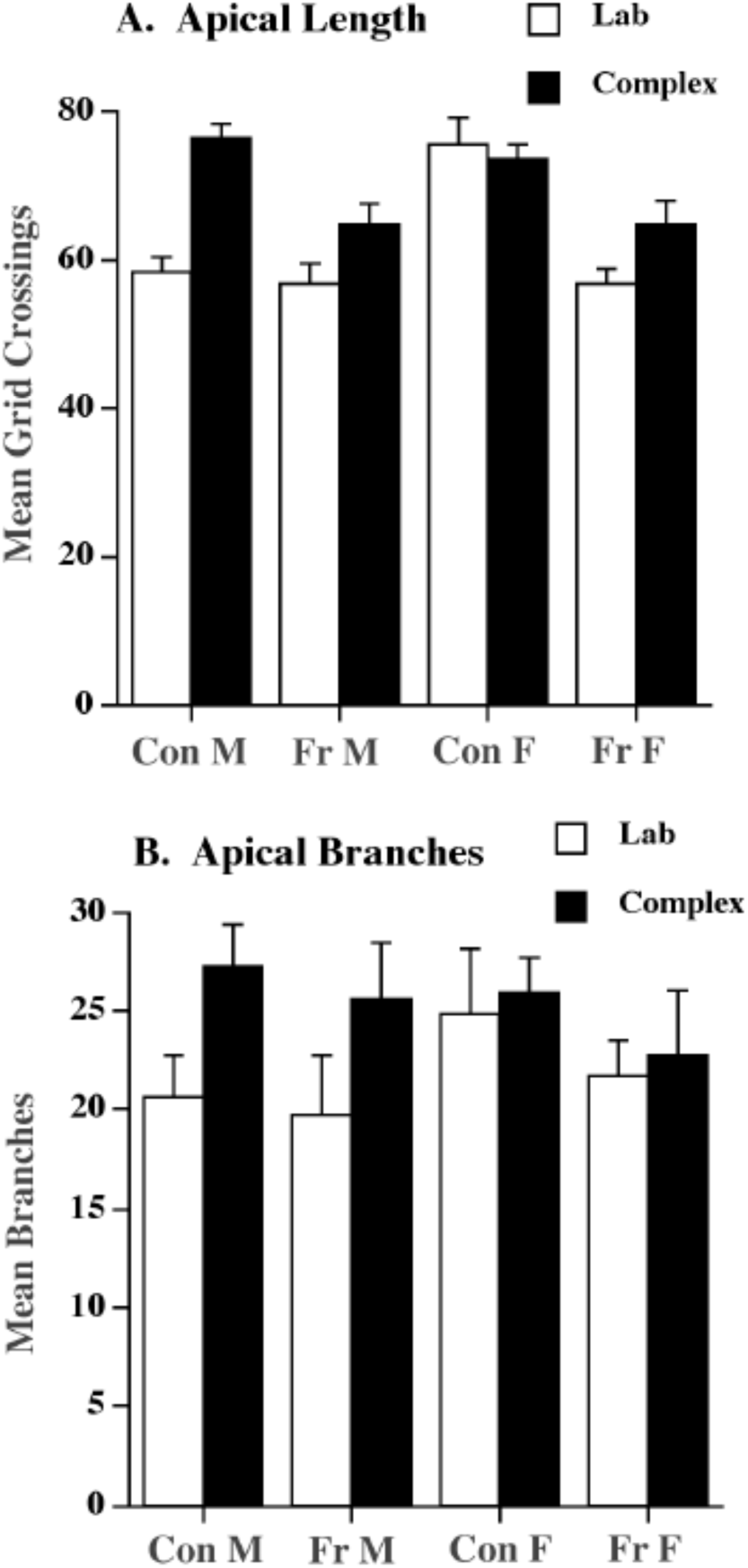
Summary of dendritic changes seen in the apical length (top) and branching (bottom) of layer III pyramidal neurons in area Occ 1. Complex housing increased dendritic length in all groups except control females. Complex housing increased dendritic branching only in males. Frontal lesions decreased dendritic length only in females.

##### Parietal Dendritic Measures

There were three overall effects. First, frontal lesions reduced dendritic length, the effect being larger in males than in females. Second, complex housing increased dendritic arborization, but the effect was larger in males than in females. Third, complex housing partly reversed the dendritic atrophy in the male, but not female, lesion animals (see Table 1).

**Table 1.**
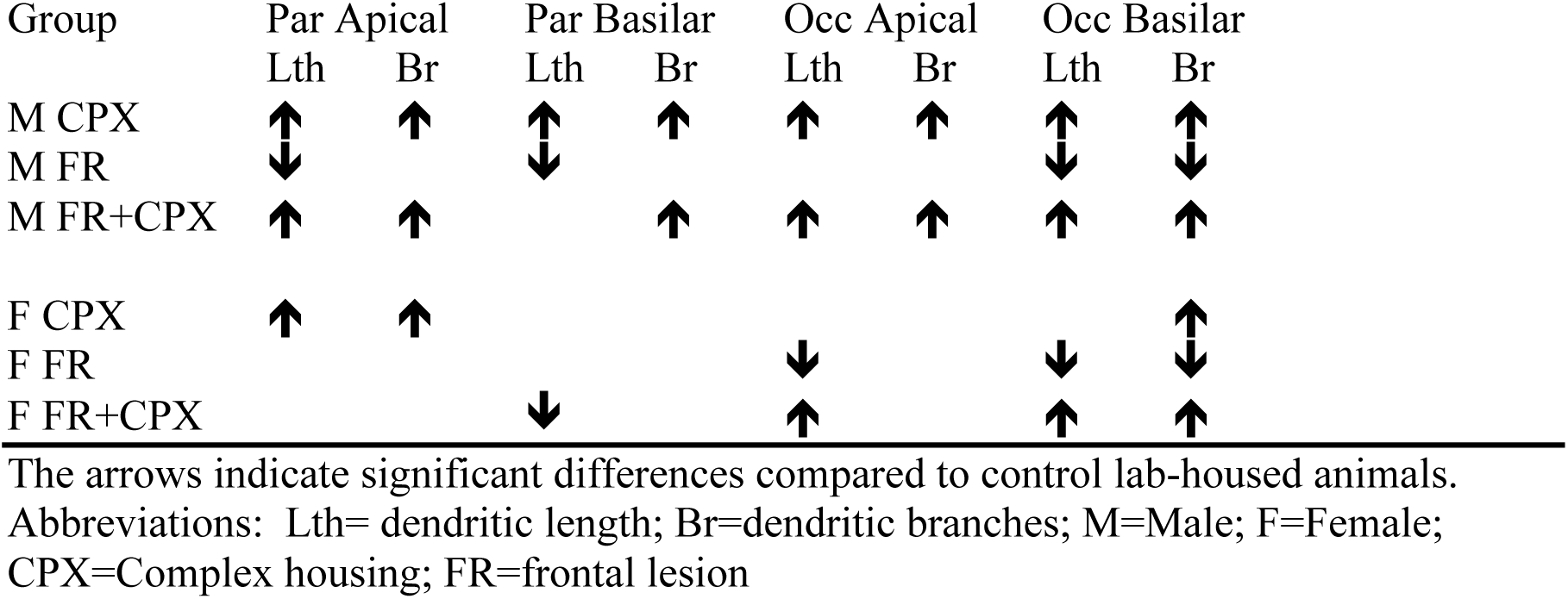
Summary of the changes in dendritic morphology.

#### Parietal dendritic length

ANOVA on the apical field found main effects of lesion (F(1,64)=5.1, p<.03) and experience (F(1,64)=17.4, p<.0001), but not sex (F(1,64)=0.001, p=.98). Only the Experience X Sex interaction approached significance (F(1,64)=3.6, p=.06), reflecting the larger experience-dependent increase in dendritic length in males than in females.

ANOVA on the basilar field found a significant effect of lesion (F(1,64)=15.5, p<.0002), but not of experience or sex (p’s>.5). Thus, both male and female frontals showed a significant drop in dendritic length. The only significant interaction was Experience X Sex (F(1,64)=4.3, p=.04), which reflected an increase in dendritic length in the male, but not the female, rats. In addition, female frontals showed a significant drop in dendritic length in response to experience.

#### Parietal dendritic branching

ANOVA on the apical field found only a main effect of Experience (F(1,64)=15.5, p<.0002) and the Lesion X Sex interaction (F(1,64)=4.7, p=.03), but no other effects were significant. Thus, all complex-housed groups showed an increase in dendritic branching. The interaction reflected the small decrease in branching in male lesion animals and a small increase in branching in the female lesion animals in both the lab and complex-housed groups.

ANOVA on the basilar field found only a Lesion X Sex interaction (F(1,64)=4.11, p<.05), again reflecting a small drop in branching in the male operates and a small increase in the female lesion animals. No other effects were significant.

#### Parietal Spine Measures

The density of spines was higher in the apical oblique segments than in the distal segments from either the apical or basilar segments. There was an increase in spine density in the male frontals but not in the female frontals. Finally, there was an overall decrease in spine density in response to complex housing, although the details varied with spine location and group (see Table 2).

**Table 2.**
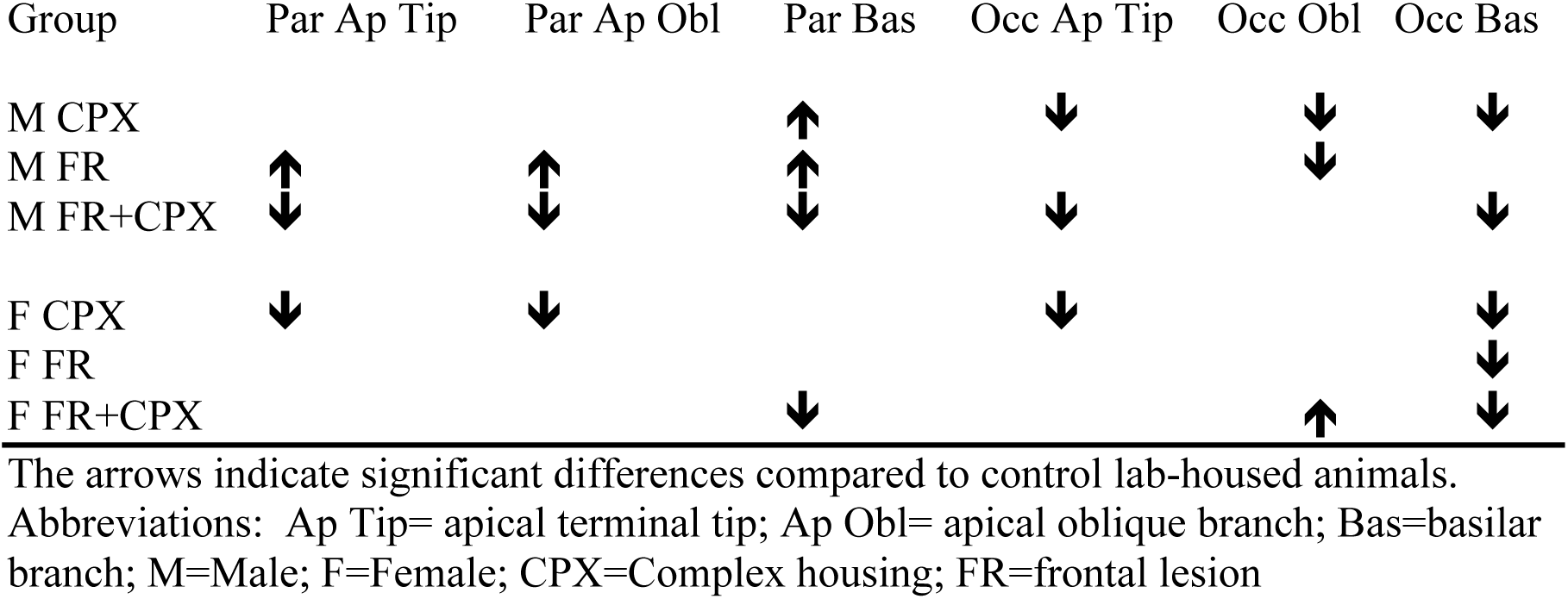
Summary of the changes in spine density.

Analysis of the apical tips for the parietal cortex revealed a significant main effect of experience (F(1,61)=10.6, p<.01), Sex (F(1,61)=5.2, p<.01), and the three-way interaction (F(1,61)=4.8, p<.03). The three-way interaction reflected the increased spine density in the frontal males, but not females, as well as the decreased spine density in response to experience. None of the other F values approached significance (p’s>.23).

ANOVA on the apical oblique segments showed a main effect of experience (F(1,61)=5.8, p<.02) and the three-way interaction (F(1,61)=3.9, p<.05. The three-way interaction reflected the decrease in spine density in response to experience in all groups except the male controls.

A similar analysis on the basilar tips found a main effect of experience (F(1,61)=6.9, p<.01), the Lesion X Experience interaction (F(1,61)=10.4, p<.002), as well as the three way interaction (F(1,61)=6.0, p<.02). The interactions reflected the fact that there was a significant increase in spine density in the frontal males and a decrease in spine density in response to experience, a result that was only significant in the lesion animals. None of the other F values approached significance (p’s>.24).

#### Occipital Dendritic Measures

Overall, the occipital measures were similar to the parietal measures with lesion-related dendritic atrophy that was reversed by complex housing (see Table 1 and Figures 8-9). *Occipital dendritic length*. Analysis of the apical fields of the occipital cortex found a main effect of lesion (F(1,64)=27.8, p<.0001), experience (F(1,64)=17.2, p<.0001), and sex (F(1,64)=3.7, p=.05). Thus, frontal lesions led to reduced dendritic length in both sexes and this was completely reversed by complex housing. All the interactions were significant (p<.05 or better) except for the Lesion X Experience interaction (p=.97). The three-way interaction reflected the fact that only the control females failed to show an effect of experience but, curiously, the control females raised in lab cages had significantly longer dendritic fields than any of the other lab-housed groups.

**Figure 9.**
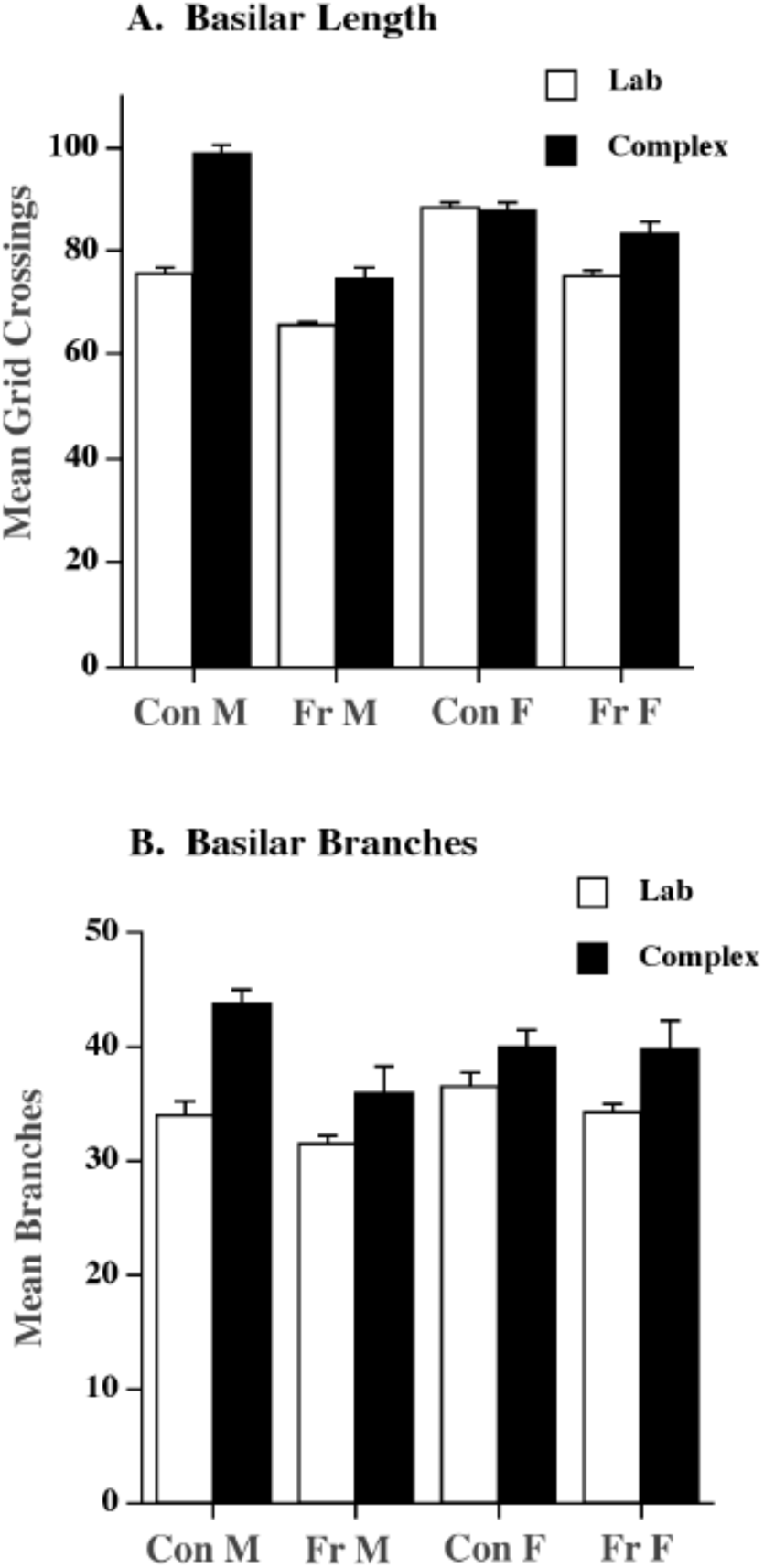
Summary of the dendritic changes seen in the basilar length (top) and branching (bottom) of layer III pyramidal neurons in area Occ 1. Complex housing increased dendritic length in all groups except control females and increased dendritic branching in all groups. Frontal lesions decreased dendritic length in both sexes.

ANOVA on the basilar fields found a main effect of lesion (F(1,64)=19.9, p<.0001) and experience (F(1,64)=11.5, p<.001), as well as the Experience X Lesion interaction (F(1,64)=4.3, p<.04) and the three-way interaction (F(1,64)=3.9, p<.05). Thus, once again the lesions led to reduced dendritic length and this was more than reversed by the complex housing experience. And, as in the apical fields, there was again no effect of experience in the females.

#### Occipital dendritic branching

The ANOVA on the apical field found main effects of lesion (F(1,64)=7.06, p<.01) and experience (F(1,64)=19.5, p<.0001) as well as an Experience X Sex interaction F(1,64)=4.3, p<.05). The reduction in dendritic branching in the lesion animals was smaller than the reduction in dendritic length but as in the length measure, it was reversed by the complex housing. The Experience X Sex interaction once again reflected the larger effect of experience in males, including no effect of experience in the control females.

Finally, ANOVA on the basilar field of the occipital cortex showed main effects of lesion (F(1,64)=8.9, p<.004) and experience (F(1,64)=30.1, p<.0001). No other effects were significant. Therefore, the lesions decreased dendritic branching, and this again was more than reversed by the complex housing.

#### Occipital Spine Measures

As in the parietal cortex, spine density was greater in the oblique branches than in the distal branches. Overall, the effect of experience was significantly reduced in the occipital cortex relative to parietal cortex, the overall finding being a *reduction* in spine density in both the apical and basilar fields (see Table 2 and Figure 10).

**Figure 10.**
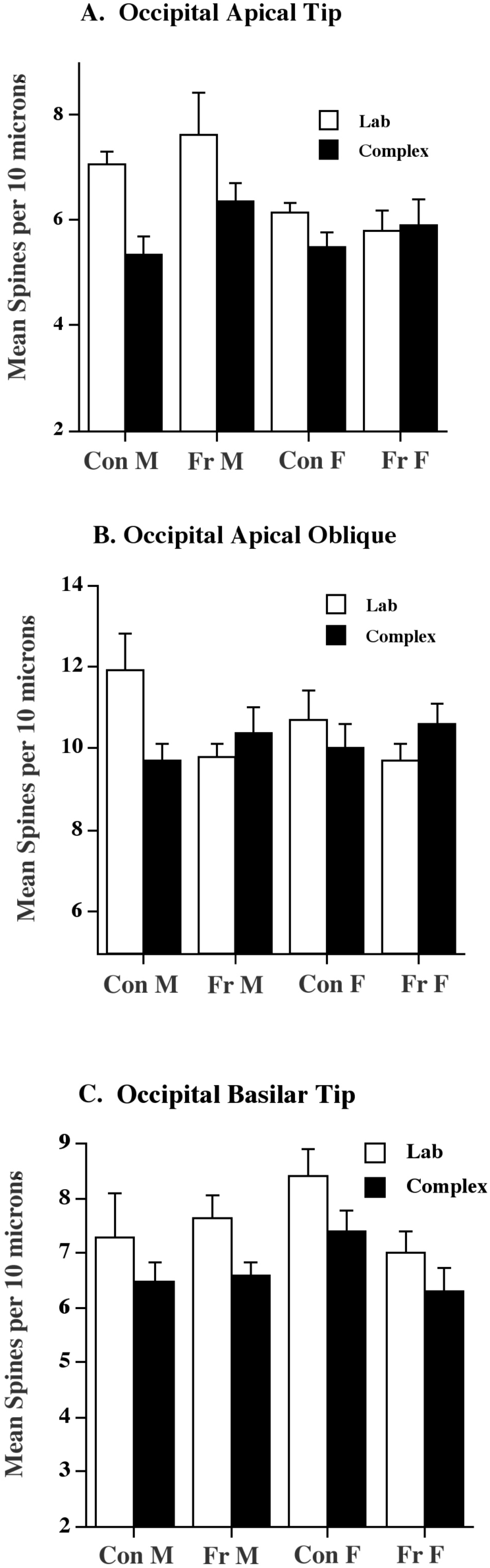
Summary of the changes in spine density in layer III neurons in occipital cortex. Frontal lesions increased spine density in the apical tip in males and reduced spine density in the basilar tip of females. Complex housing generally reduced spine density in males but less so in females.

#### Occipital spines

ANOVA on the apical tip found a main effect of experience (F(1,52)=9.2, p<.004), sex (F(1,52)=8.7, p=.005), and the Experience X Sex interaction (F(1,52)=4.6, p<.04). Thus, experience generally decreased spine density but the effect was larger in males and absent in the female operates. None of the other F values approached significance (p’s>.20).

ANOVA on the oblique segment showed an effect only on the Lesion X Experience interaction (F(1,52)=6.9, p=.01). This result reflected a single effect of reduced spine density in the control males housed in complex environments.

Finally, ANOVA on the basilar tip found a main effect of experience (F(1,52)=7.6, p=.008) and the Lesion X Sex interaction (F(1,52)=5.1, p<.03). The interaction reflected that finding that whereas males with frontal lesions showed an increase in spine density, females showed a significant decrease. Again, none of the other F values approached significance (p’s>.13).

## DISCUSSION

There three novel findings in these studies. First, there were sexually-dimorphic effects of experience on functional recovery from P7 frontal injuries. Second, the early lesions produced a sexually-dimorphic atrophy of cortical pyramidal cells in both the parietal and occipital cortex, the effect being larger in males in parietal cortex and larger in females in occipital cortex. Third, complex-housing rats with early frontal lesions reversed the lesion-induced changes in dendritic arborization, the effect being larger in males than females. In addition, we confirmed earlier studies of the effect of day 7 frontal lesions on behavioral and cortical development. We consider each main finding separately.

### Complex housing has sex-specific effects on recovery from day 7 frontal lesions

The results of the current study extend our previous results showing that day 7 medial frontal cortical lesions have sexually dimorphic effects on both behavior and anatomy in lab-reared animals (Kolb and Stewart, 1995; Kolb et al., 1996). In our earlier studies we found that male rats with day 7 medial frontal lesions showed better performance on spatial learning tasks than did female rats with similar lesions. The current study showed that although this was once again true, females showed better spontaneous recovery on a test of skilled motor behavior and in contrast to males, showed some benefit of complex housing. We did not examine skilled reaching (or any other motor behaviors) in our previous studies looking at the effects of complex housing.

The unexpected sexual-dimorphism on different measures of functional recovery is instructive and suggests that studies claiming sex differences in functional recovery from cerebral injury at any age will need to examine a wider range of behaviors than is normally done. The medial prefrontal cortex has an important role in the control of skilled forelimb movements, presumably because of the direct corticospinal connections from the anterior cingulate cortex, as well as the better-studied effects on cognitive behaviors.

### Complex housing has sex-dependent effects on neuronal morphology

The current study replicates findings shown previously by many investigators who have shown that housing animals in various types of complex environments increases dendritic arborization in cortical neurons (e.g., Coleman and Riesen, 1968; Rosenzweig and Bennett, 1978; Sirevaag and Greenough, 1988; Uylings, Kuypers, Diamond, & Veltman, 1978; Walsh, 1982). Furthermore, the current study replicates the finding of Juraska and her colleagues (e.g., Juraska, 1984; 1990) that there is a basic sex difference in the effect of complex housing, at least in visual cortex. Thus, whereas males show an increase in dendritic arborization across the cortex, females show a reduced effect in sensorimotor cortex and virtually no effect at all in occipital cortex (see Table 1).

The current study failed to replicate our finding that placing male control animals in complex environments at weaning leads to a *decrease* in spine density in the parietal cortex. There was no effect of experience on the apical field. Females did show the decrease in spine density in parietal cortex, however. What was novel was that both sexes showed decreased spine density in occipital cortex. This decrease (or no change) in spine density contrasts to the effects of placing animals in complex environments later in life, which leads to an *increase* in spine density (e.g., Kolb et al., 2003; Globus et al., 1973). It thus appears that similar experiences can have qualitatively different effects at different ages. In particular, it appears that sensory and/or motor stimulation has unexpected effects on the young brain. In fact, it is not just the weanling brain that shows these effects. In particular, we have found that newborn rats that are tactilely-stimulated daily with a small paintbrush over the first two weeks of life also show a decrease in spine density when the brains are examined in adulthood (Kolb & Gibb, 2010).

Our finding of a decrease in spine density in young animals has precedents (e.g., Bock and Braun, 1998; Wallhausser and Scheich, 1987). For example, Wallhausser and Scheich (1987) presented newly hatched chicks with a hen or an acoustic stimulus with the goal of imprinting the chicks to the visual or auditory stimulus they found that neurons in the hyperstriatum of the imprinted chicks showed decreased spine density in the trained chicks when the brains were examined 7 days after training. There is an important caveat to this result, however. Patel, Rose & Stewart (1988) used a Golgi technique to impregnate chick brains 25 hours after training chicks to avoid a colored bead. They found a twofold *increase* in spine density in the neurons in a region of the hyperstriatum in the “trained” chicks. The critical difference between the Wallhausser and Scheich study and the Patel et al study is the survival time. Thus, taken together the experiments show that there was an increase 25 hours after training but there was a decrease 7 days after training. The simplest conclusion from the chick studies is that the novel stimulation may cause an initial rapid increase in spine density, followed by a pruning. If we extrapolate to the current study, we might predict that the juvenile animals showed an increase in spine density over the first hours or days in the complex environments, followed by a synaptic pruning.

At this point we do not know what the drop in spine density in the young animals actually reflects. It is possible that the early experience also stimulates changes in glial proliferation or in neuronal apoptosis. In the latter case, it could therefore be that there are more neurons, but each neuron has fewer connections. This remains to be proven, however.

### Neonatal frontal lesions have sex-dependent effects on neuronal morphology

The results of the current study show that P7 frontal cortical lesions have sexually dimorphic effects on neuronal morphology. Thus, following day 7 medial frontal lesions males show an increase in spine density in pyramidal neurons in parietal cortex that is not observed in females. This increased spine density may play some role in facilitating recovery on measures of cognitive behavior, such as the spatial learning measure in the current study, but it evidently does not facilitate recovery of skilled motor behavior. At this point we do not know what morphological changes might correlate with the improved motor skill performance in the females, although we can propose that there may be lesion-related differences in morphology of forelimb neurons in motor cortex. It was not practical to draw motor cortex neurons in the current study, however, because we have shown elsewhere that there is an idiosyncratic shift in location of the motor maps of animals with neonatal frontal lesions (Williams et al., 2006). Given that the location of this map shift cannot be determined from Golgi-stained tissue, we thus would be unable to be certain that we were measuring functional similar neurons in different animals. The critical study would be one in which the animals had motor maps identified by electrical stimulation, followed by Golgi analysis.

The failure to find experience-dependent benefits on the spatial learning task in female frontals may be related to sex differences in the effect of both injury and experience on the occipital neurons. Occipital neurons show increased dendritic length when animals are trained on visual spatial tasks (e.g., Greenough & Chang, 1989; Kolb et al., 2008), which suggests that changes in occipital neurons may be important in spatial learning. Although animals with frontal injuries in both sexes showed a decrease in dendritic length in occipital neurons, relative to same-sex control animals, the decrease was larger in females.

One explanation for the sex-related differences in both the response to injury and to experience may be related to sex differences in epigenetic responses to these experiences. Although we have not specifically looked at sex differences in epigenetic measures in response to complex housing and injury, there is a growing both of evidence that there are sex differences in response to early life experiences and brain injury (for a review, see Arambula et al., 2019). In addition, we have preliminary evidence that there are sex differences in both global methylation as well as gene expression in cortex following early stress in rats (Mychasiuk et al., 2011a; 2011b). It is reasonable to predict that these injury-related sex differences will lead to differential effects of experience in males and females. The challenge will be to relate such epigenetic differences directly to neuronal morphology and function.

## CONCLUSION

Most preclinical studies of the beneficial effects of complex housing after brain injury have focused on cerebral injuries in adults (for extensive reviews, see Alwis & Rajan, 2014; McDonald et al., 2018). Overall, complex housing generally has a larger effect on cognitive functions than on complex motor functions, although complex housing, in combination with other interventions, does show promising effects on skilled motor functions (e.g., Corbett et al., 2015; Jeffers & Corbett, 2018). There are fewer studies of the benefits of complex housing following brain injury in infancy, but the effects appear to be similar with improved cognitive functions and only small, if any, effects on complex skilled motor functions. We have, however, found beneficial effects of tactile stimulation in rats with either medial frontal lesions (Kolb & Gibb, 2010) or hemidecortications (Day et al., 2023), although the lesion animals still show significant impairments. As with the Corbett studies in adult stroke animals it might be prudent to combine both complex housing and tactile stimulation in animals with neonatal cortical lesions. Indeed, given that the application of complex housing experiences in human clinical trials have used multiple experiences such as computer games, music, novel training tasks etc (Alwis & Rajan, 2014; McDonald et al., 2018), it would be reasonable to try to use multiple treatments in the preclinical studies as well, with special emphasis on motor tasks.

